# ADSoluble aggregates present in cerebrospinal fluid change in size and mechanism of toxicity during Alzheimer’s disease progression

**DOI:** 10.1101/600346

**Authors:** Suman De, Daniel R. Whiten, Francesco S. Ruggeri, Craig Hughes, Margarida Rodrigues, Dimitrios I. Sideris, Christopher G. Taylor, Francesco A. Aprile, Serge Muyldermans, Tuomas P. J. Knowles, Michele Vendruscolo, Clare Bryant, Kaj Blennow, Ingmar Skoog, Silke Kern, Henrik Zetterberg, David Klenerman

**Affiliations:** Department of Chemistry, University of Cambridge, Cambridge CB2 1EW, United Kingdom; Centre for Misfolding Diseases, University of Cambridge, Cambridge CB2 1EW, United Kingdom; Department of Veterinary Medicine, University of Cambridge, Cambridge, CB3 0ES, United Kingdom; Laboratory of Cellular and Molecular Immunology, Vrije Universiteit Brussel, Brussels, Belgium; Clinical Neurochemistry Laboratory, Unit of Department of Psychiatry and Neurochemistry, Institute of Neuroscience and Physiology, the Sahlgrenska Academy at the University of Gothenburg, Sweden; Clinical Neurochemistry Laboratory, Sahlgrenska University Hospital, Mölndal, Sweden; Neuropsychiatric Epidemiology Unit, Department of Psychiatry and Neurochemistry, Institute of Neuroscience and Physiology, the Sahlgrenska Academy at the University of Gothenburg, Sweden; Department of Neurodegenerative Disease, UCL Queen Square Institute of Neurology, University College London, Queen Square, London, UK; UK Dementia Research Institute at University College London, London, UK; UK Dementia Research Institute at University of Cambridge, Cambridge CB2 0XY, United Kingdom

## Abstract

Soluble aggregates of amyloid-β (Aβ) have been associated with neuronal and synaptic loss in Alzheimer’s disease (AD). However, despite significant recent progress, the mechanisms by which these aggregated species contribute to disease progression are not fully determined. As the analysis of human cerebrospinal fluid (CSF) provides an accessible window into the molecular changes associated with the disease progression, we studied the soluble Aβ aggregates present in CSF samples from individuals with AD, mild cognitive impairment (MCI) and healthy controls. We found that these aggregates vary structurally and in their mechanisms of toxicity. More small aggregates of Aβ that can cause membrane permeabilization already found in MCI; in established AD, the aggregates were larger and more prone to elicit a pro-inflammatory response in glial cells. These results suggest that different neurotoxic mechanisms are prevalent at different stages of AD.

## Introduction

Small soluble aggregates of amyloid-β (Aβ) have been shown to impair hippocampal synaptic plasticity, induce learning deficits and correlate with cognitive impairments both in Alzheimer’s disease (AD) mouse models and humans^1–4^. There soluble aggregates can exert cellular toxicity via a range of diverse mechanisms, including oxidative stress, disruption of Ca^2+^ homeostasis and cellular signalling, mitochondrial alterations, glial activation and inflammation^5,6^. Many of these processes are the consequence of two fundamental upstream events induced by soluble aggregates: *(i)* permeabilisation of cell membranes by non-specific binding^7,8^ and *(ii)* specific interactions with receptors in cell membranes^6,9^. It is also known that the morphology of protein aggregates can determine the level of their involvement in different biological interactions. Hydrophobic protein aggregates more readily interact with hydrophobic lipid membranes, while the size and shape of the aggregates determine the affinity of binding to receptors^6,8–10^. Understanding how the intrinsic heterogeneity in the size, shape and structure of the aggregates present in the human brain influences their mechanism of toxicity and how such heterogeneity changes as AD progresses is crucial in identifying the molecular pathways that lead to neuronal death.

Most of our current knowledge about the origins and morphologies of the toxic soluble species involved in neurodegenerative mechanisms is derived from *in vitro* studies and animal models. These studies have shown the presence of soluble aggregates with large heterogeneity in size (dimer to higher order multimers), shape (small spherical to fibril like) and structure (random coil to β-sheet)^4,5^. During disease progression, Aβ in the human brain can aggregate into a large number of different forms, consisting of different numbers of peptides, sizes, shapes and structural configurations, with multiple possible post-translational modifications and co-aggregating species. One way to characterise the presence of toxic aggregates at different clinical stages and its implication on disease progression is to study these soluble aggregates present in AD, mild cognitive impairment (MCI) and healthy control cerebrospinal fluid (CSF), since this fluid can reflect at least some of the biochemical changes occurring inside the brain. MCI is a heterogeneous syndrome, which may have many underlying causes^11^. A proportion of MCI patients have AD pathology, and are at increased risk of developing AD dementia. MCI can thus be conceptualised as a prodromal state in the AD continuum - a transition between normal cognitive aging and AD^11,12^. In this study, we used the core AD CSF biomarkers and critical dementia rating (CDR), which depend on the cognitive ability of individuals (see Methods), to distinguish the MCI cases that had Alzheimer’s pathologic changes. We have selected MCI cases based on the low level of Aβ42 (Aβ42 < 600 ng/L) indicating brain amyloidosis and where the CDR is at least 0.5. This procedure allowed us to select solely for the MCI cases with brain amyloidosis^13^, which will be referred to from now on as MCI. All the healthy controls are free of MCI and dementia and had a CDR of 0. Therefore, to understand the nature of soluble aggregates present at different stages of AD and how they induce cellular toxicity, we studied CSF samples collected from individuals affected from AD and MCI and compared these with healthy controls.

CSF is continuously produced, recycled and freely exchanged with the interstitial fluid in the brain, making it an ideal reservoir of soluble aggregates that can be reflective of toxic species present in the brain tissue for that disease stage. It has been demonstrated that soluble Aβ is secreted into the CSF, making it an ideal biomarker candidate for AD^11,14,15^. Although the concentrations of Aβ oligomers and monomers have been measured in AD CSF, due to a lack of suitably sensitive methods, the specific detection and quantification of the heterogeneity in morphology of soluble Aβ aggregates present in CSF have not yet been determined. Understanding the changes in these soluble aggregates that occur during disease progression may provide new insights into the disease mechanisms with potential for early diagnosis.

### Soluble aggregates present in AD and MCI CSF induce toxicity by distinct mechanisms

To achieve this goal, we set out to analyse the toxicity of the soluble aggregates in CSF by measuring their ability to permeabilise lipid membranes and induce an inflammatory response. We used CSF from individuals diagnosed with MCI, AD as well as healthy controls. As the unspecific binding of protein aggregates to lipid membranes is driven in particular by hydrophobic interactions, aggregates with more hydrophobic patches show increased toxicity via disruption and permeabilisation of the lipid membrane. To quantitatively measure how the soluble aggregates act in this manner, we used a recently developed biophysical assay, which has shown that physiological concentrations of soluble aggregates of Aβ can destabilise lipid membranes^16,17^. For this assay, we immobilised thousands of single POPC vesicles (mean diameter - 200 nm) containing the Ca^2+^-specific dye, Cal-520, onto PEGylated glass cover slides via biotin-neutravidin linkage. When CSF samples are added, soluble aggregates present permeabilise the membrane of lipid vesicles and Ca^2+^ from the surrounding solution enter individual vesicles causing a change in the fluorescence intensity of the dye (**Fig. 1a**). The change in fluorescence intensity in these nano-sized vesicles is proportional to the number of ions that entered and can be quantified using TIRF microscopy^16,18^. Using this method, we found that aliquots of MCI CSF cause greater membrane permeabilisation compared to the AD and control CSF (**Fig. 1b**). By contrast, we found no significant difference in membrane permeation induced by AD and control CSF, in agreement with a previously reported study^19^.

**Figure 1.**
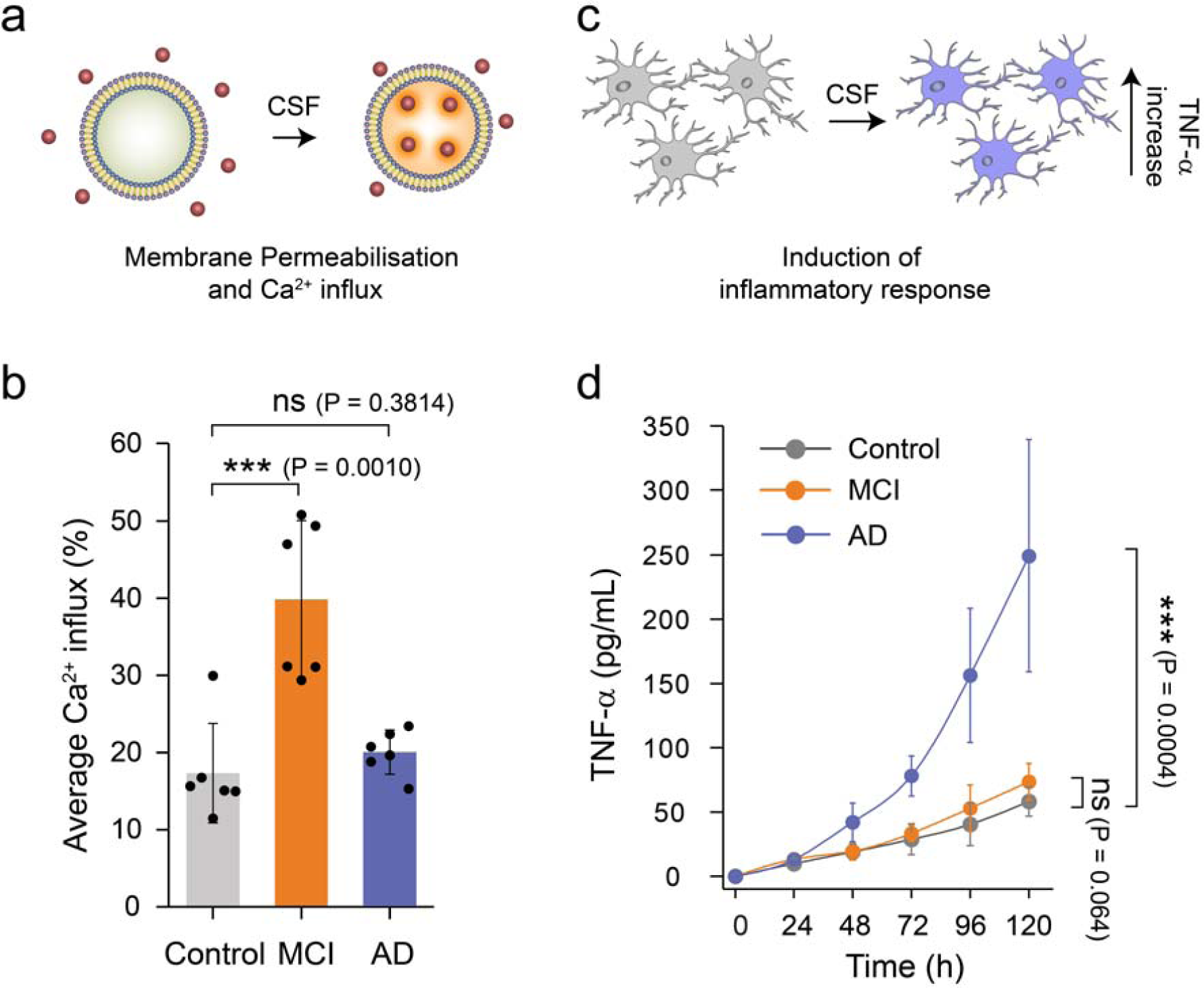
Soluble aggregates present in MCI and AD CSF samples show different dominant mechanisms of toxicity. **(a)** Membrane permeabilisation was measured by immobilising hundreds of vesicles containing a Ca^2+^-sensitive dye on PEGylated glass cover slides. If any species present in the CSF disrupts the integrity of the lipid membrane of the vesicles, Ca^2+^ ions from the surrounding buffer enter into individual vesicles in numbers that can be quantified using highly sensitive TIRF microscopy. **(b)** Aliquots of MCI CSF can cause more membrane permeabilisation compared to AD and control CSF (n= 6 AD, 6 MCI, 6 control CSF). A two-sample unpaired t-test was performed to compare each data set. **(c)** The inflammatory response in microglia cells was quantified using an ELISA assay to measure the levels of secreted tumour necrosis factor alpha (TNF-□). **(d)** For this study, CSF samples were added to BV2 microglia cells and incubated for 120 h. Every 24 h the TNF-α concentration in the supernatant was quantified using an ELISA assay. AD CSF samples were more effective MCI and control CSF samples in inducing an inflammatory response (lines are guide to the eye; n= 10 AD, 6 MCI, 6 control CSF). Error bars are the standard deviation among data points. A two-sample unpaired t-test at the 120h time point was performed to compare the data sets.

We instead observed the opposite trend in the CSF-induced pro-inflammatory response in glial cells (**Fig. 1c**). Physiological concentrations of protein aggregates can interact with specific membrane receptors in microglial cells and induce a proinflammatory response^20^. This response can be quantified by measuring secreted tumour necrosis factor alpha (TNF-□), one of the pro-inflammatory cytokines that is produced, using an ELISA assay. To determine if CSF samples can induce a proinflammatory response, we added them to BV2 microglia cells and incubated for 5 days. Every 24 h, we took the supernatant above the cells and measured the TNF-α concentration. We found that CSF aliquots can activate an innate immune response through the production of significant amounts of TNF-α. We observed that AD CSF samples induced a stronger inflammatory response than the MCI and control CSF samples (**Fig. 1d**). This inflammatory response induced by AD CSF was significantly blocked by *Rhodobacter sphaeroides* lipid A (RSLA), a known Toll-like receptor (TLR)-4 antagonist^21^ and TAK-242, a small-molecule inhibitor that binds selectively and inhibits signalling of TLR-4^22^ (**Supplementary Fig. 1**).

### Soluble amyloid-beta aggregates present in AD and MCI CSF are structurally distinct

To gain insight into the composition of the soluble aggregates responsible for the observed membrane permeabilisation, we employed an Aβ-specific antibody (Nb3) previously shown to counteract the toxicity induced by soluble aggregates of full length as well as N-terminally truncated Aβ both *in vitro*^16,23^ and AD CSF^19^. We found that Nb3, which recognises amino acids 17-28 of the Aβ sequence, was able to significantly reduce the aggregate-induced membrane permeabilisation for both AD and MCI (**Fig. 2a and 2b**). This result indicates that at least a portion of the toxic aggregates present in MCI and AD CSF are composed of Aβ. We also found that an antibody that targets the N-terminus of Aβ (3-9), but not an antibody that targets the C-terminus (36-42)^24^, efficiently inhibits AD CSF-induced inflammatory response (**Fig. 2c**). The differential effects of N-terminal and C-terminal antibodies provide information not only on the composition but also on the structure of the soluble aggregates present in AD CSF. Mature fibrillar aggregates have a hydrophobic C-terminus which is inaccessible to C-terminal antibodies, while the N-terminus residues of aggregates are exposed^25–27^. However, the N-terminus and C-terminus antibodies were both equally effective at reducing lipid membrane permeation (**Fig. 2d**), suggesting that the aggregates present in the CSF that are responsible for inducing membrane permeation are structurally distinct from those that induce an inflammatory response.

**Figure 2.**
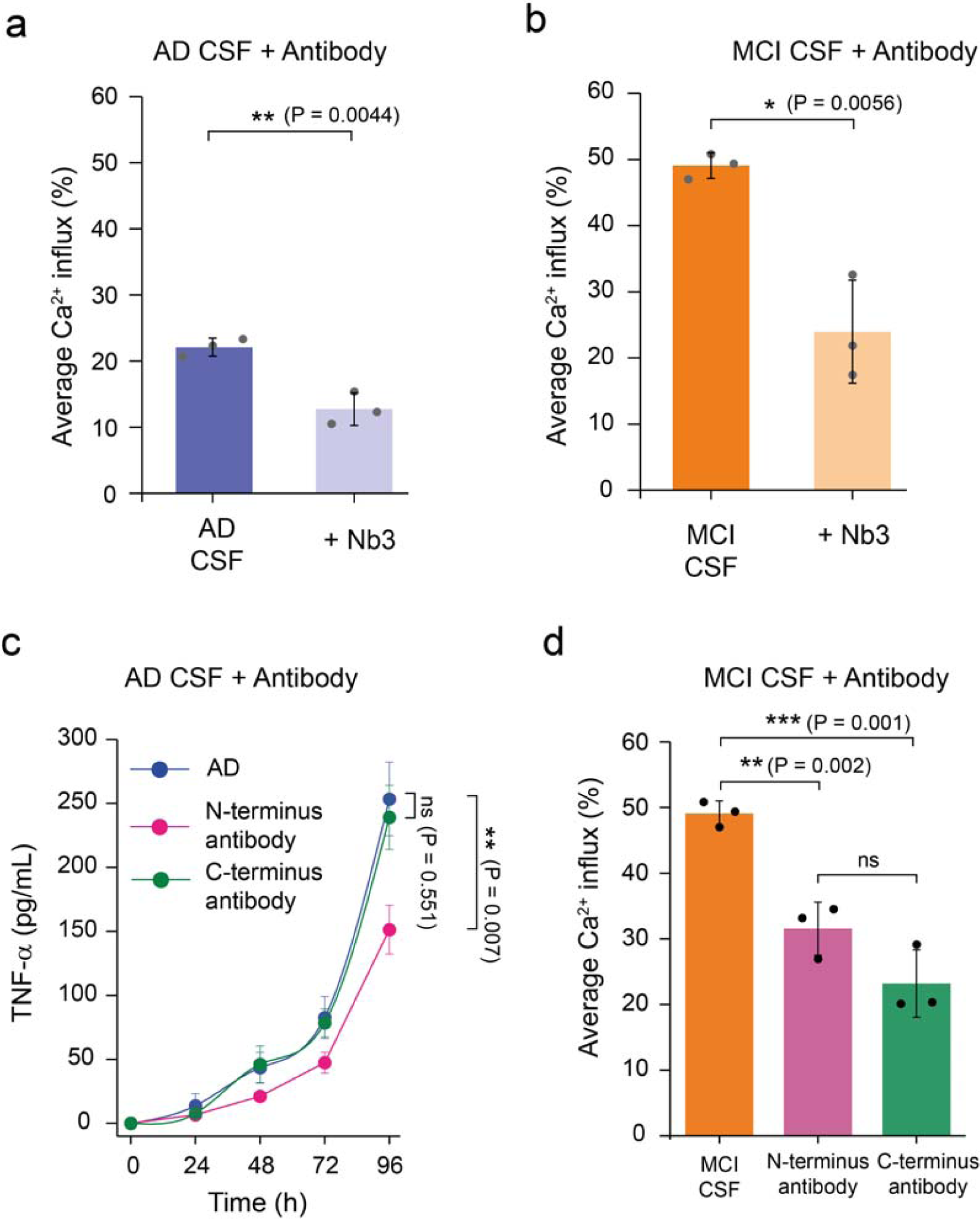
The toxic soluble aggregates present in MCI and AD CSF contain **A**β. A significant inhibition of the membrane permeabilisation of lipid membranes by **(a)** AD and **(b)** MCI CSF samples is caused by a nanobody Nb3 (300 nM) designed to bind to Aβ, indicating that some of the aggregates present in MCI and AD CSF contain Aβ. Nb3 recognises amino acids 17-28 of the Aβ sequence and shown to inhibit toxicity induced by protein aggregates of Aβ cerebrospinal fluid samples from AD patient **(c)** A N-terminal binding antibody (binding the region of residues 3-9 of Aβ42) is more potent at reducing the aggregate-induced inflammatory response than a C-terminal designed antibody (binding the region of residues 36-42 of Aβ42); P values are calculated using a two sample t-test to compare the inhibition by an N-terminally binding antibody and a C-terminally antibody at 96 hours (n=3). Error bars are the standard deviation among data points. **(d)** Both N-terminally and C-terminally targeting antibodies significantly inhibit MCI CSF-induced membrane permeabilisation. However, we do not find any significant difference between their activities. Error bars are the standard deviation among data points. Two sample unpaired t-tests were performed to compare the data sets (n= 3).

### Single aggregate based super-resolution imaging shows that the size distribution of the soluble aggregates present in AD and MCI CSF are different

To examine the size and morphology of these soluble aggregates, we employed a newly developed super-resolution technique, aptamer DNA PAINT (ADPAINT) (**Fig. 3a**)^28^. The small size of the aptamer in combination with its remarkable specificity and affinity allows us to resolve aggregated structures of Aβ with a precision of about 20 nm. We have previously demonstrated the utility of this imaging technique by accurately identifying and nanoscopically characterising the shapes and sizes of the species formed during an aggregation reaction and in fixed neurons^28^. Using this technique, we analysed the average number of aggregates present in the CSF samples and measured the size of individual aggregates (**Fig. 3b,c**), which ranged from 20 nm, limited by our resolution, to 300 nm. We did not find any significant difference in the total number of aggregated species present among MCI, AD and control CSF samples (6 each), a result that we confirmed using the amyloid-specific dye, pentameric formylthiophene acetic acid (PFTAA) (**Supplementary Fig. 2**).

**Figure 3:**
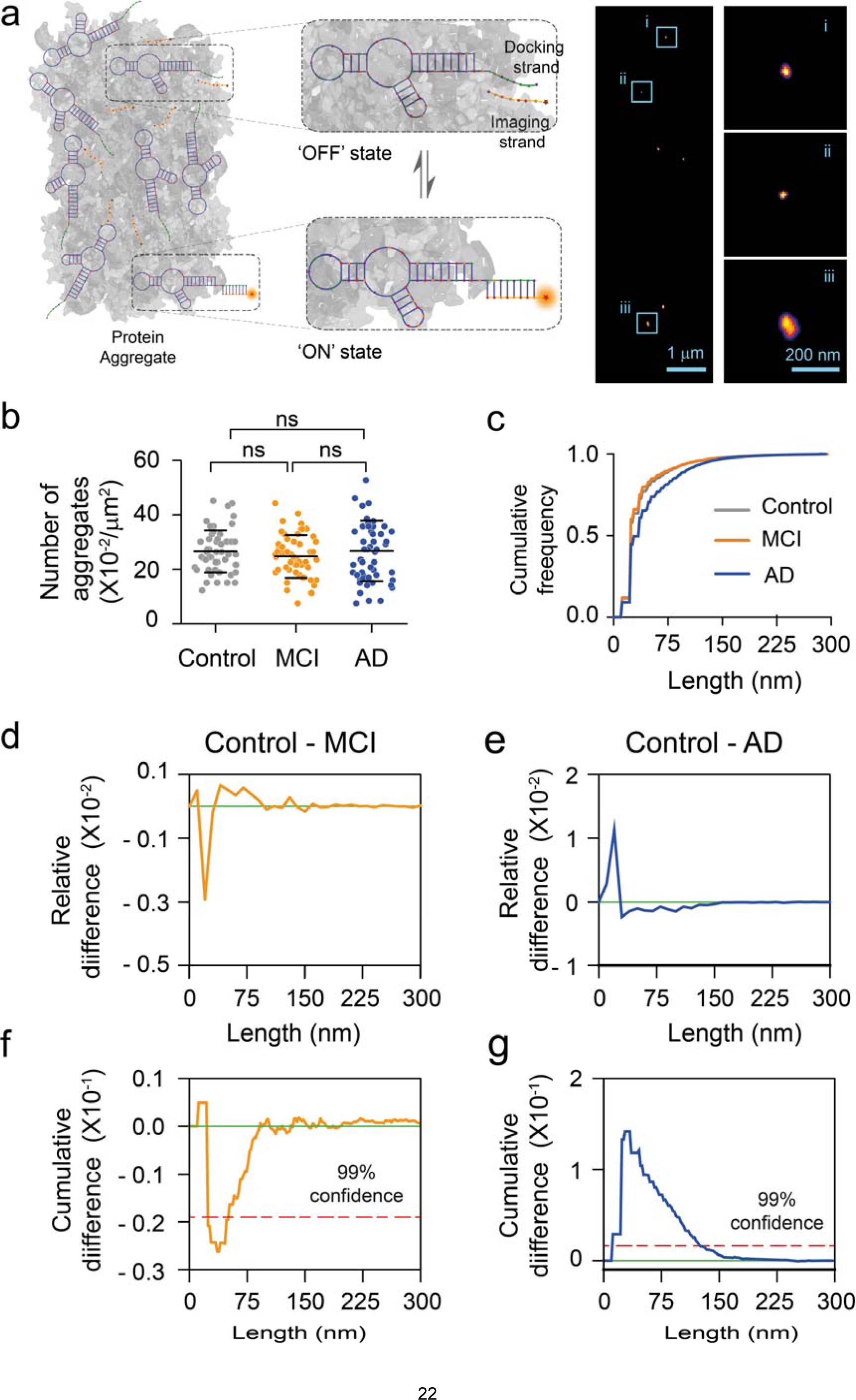
Super-resolution imaging of aggregates present in CSF using AD-PAINT. (a) Schematic of AD PAINT. Dye-labelled DNA imaging strands transiently bind to their complementary target sequence (docking strand), which is attached to a protein aggregate via aptamer. This transient binding between imaging and docking strand is detected. Repeated cycles of binding and unbinding allows a super-resolved image of individual protein aggregate present in CSF to be determined. The right image shows examples of super-resolved image of protein aggregates from CSF. Three individual protein aggregates present in CSF are enlarged. The lengths of the aggregates shown are: (i) 47 nm, (ii) 34 nm and (iii) 118 nm. (b) Number of aggregates present in control, MCI and AD CSF samples. Each point represents one field of view. (c) Cumulative frequency histograms of the size distributions of all aggregates measured. (d,e) Differences between normalised histograms of the size distributions of the indicated CSF samples. (f,g) Differences between the cumulative frequency histograms of the size distributions of the indicated CSF samples. The dotted line indicates 99% confidence using the Kolmogorov-Smirnov statistical test. (n= 6 AD, 6 MCI, 6 control CSF).

To examine the aggregate morphologies, we plotted the relative differences between normalised (**Fig. 3d,e**) and cumulative (**Fig. 3f,g**) histograms of the size distributions of aggregates present in control vs MCI and AD CSF samples. In the plots of relative differences negative values indicate that more aggregates of that size are present in the disease CSF, whereas positive values indicate more aggregates of that size are present in control CSF. Similarly, a negative slope indicates more aggregates of that size are present in disease CSF when looking at the cumulative differences. We found that, with 99% confidence, the number of small aggregates (< 50 nm) present in MCI CSF was higher than that found in control CSF (**Fig. 3d,f**). This result suggests that a relatively larger number of small aggregates might be accountable for enhanced membrane permeabilisation induced by MCI CSF. This finding agrees with previously reported studies that small sized Aβ aggregates are more hydrophobic and have greater tendencies to interact, permeabilise and cross the plasma membrane^8,10,29^. In contrast, a ten-fold larger change was observed in the size distribution of AD CSF compared to control CSF, with a larger number of longer, mature aggregates (∼ 40 to 200 nm) (**Fig. 3e,g**). This finding suggests that relatively longer aggregates present in AD CSF might be responsible for the increased inflammatory response. These results, in combination with AD CSF induced inflammation inhibition by N-terminus antibody (**Fig. 2a**), are also in agreement with previous reports. It has been shown that Aβ aggregates greater than 100 nm formed in artificial CSF can trigger an inflammatory response in microglial cells that can be blocked by an Aβ N-terminal region recognising antibody^27,30,31^.

### Structural characterisation of soluble aggregates presents in CSF using high resolution atomic force microscopy at the single aggregate level

To check the heterogeneity and three-dimensional morphological properties of the protein aggregates present in the CSF samples, we utilised a phase controlled atomic force microscopy (AFM) technique to resolve the structures of individual protein aggregates at Angstrom resolution ^32,33^ (**Fig. 4**). The control CSF sample showed the uniform presence of abundant spherical species, whereas both the MCI and AD samples showed the co-existence of spherical species and elongated aggregates (**Fig. 4**). To unravel the heterogeneity and structural properties of the protein aggregates in the MCI and AD CSF samples, we performed a statistical analysis of their cross-sectional length and heights of individual aggregates^33^. The MCI CSF sample showed a relatively uniform population of elongated aggregates with a cross-sectional height ranging between 0.3-1 nm (**Fig. 4b**) and length 50-100 nm (**Fig. 4c**), larger than the typical length of species in the control sample (p<0.001). The AD CSF samples showed the coexistence of highly heterogeneous elongated aggregates. These species belonged roughly to two populations of height, the first with cross sectional height ranging between 0.3-1 nm which is similar to the species present in MCI CSF aggregates, often referred to as protofilaments, and the second with cross-sectional height between 1-3 nm, often referred to as protofibrils. A larger cross-sectional aggregate height has been associated *in vitro* to an increased content of intermolecular β-sheet and maturity of amyloid structure^32^. The AD elongated aggregates are significantly longer than the MCI CSF sample (p<0.001), ranging in length between 50-400 nm. These data, in conjunction with the AD PAINT data, confirm that the protein aggregates present in AD CSF differ in structure from the aggregates present in MCI and control CSF. Interestingly, there is a crystal structure of TLR3, which is in the same family as TLR4, bound to an RNA dimer which is about 2 nm in diameter^34^. TLR3 signalling occurs when the RNA dimer is longer than 15 nm^35^. This observation suggests that the long AD protofibrillar aggregates, with cross-sectional height of round 2 nm, might be the species responsible for TLR4 signalling in AD CSF.

**Figure 4.**
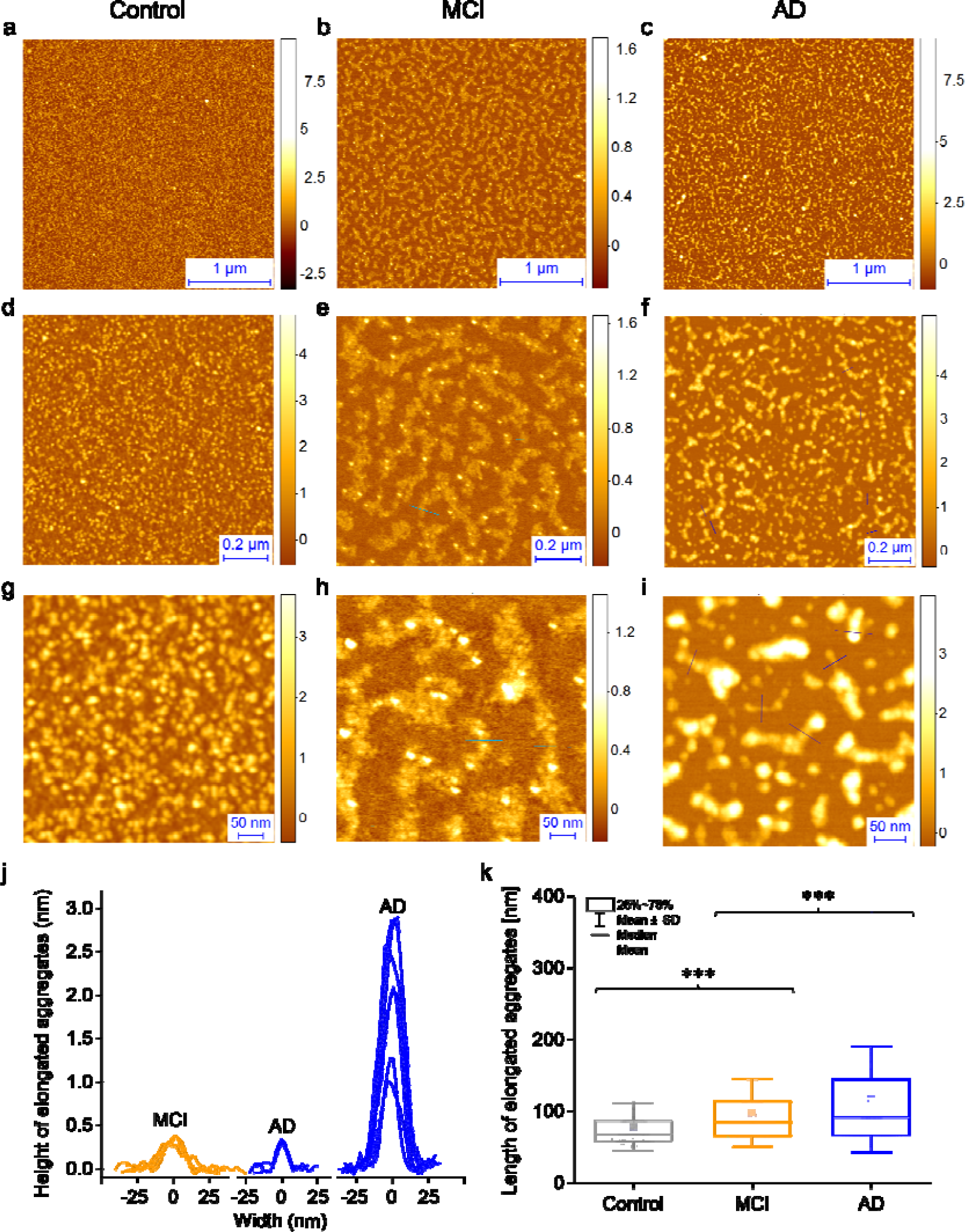
Characterization of the protein aggregates present in CSF samples at single aggregate level using AFM. (a-c) Representative AFM images of the aggregates present at different CSF samples with (d-j) a magnified example of the species present. (j,k) Statistical analysis of the cross-sectional length and heights of individual aggregates present in CSF samples; the length of the individual aggregates are shown as a box chart.

### Conclusions

In conclusion, we have reported that there are more number of small aggregates that can cause increased membrane permeabilisation present in MCI CSF than in AD CSF, while in AD CSF there is an increased number of larger and longer aggregates, corresponding with enhanced neuroinflammation. These larger aggregates are protofibrils, with distinct structural features from those of the smaller aggregates and appear to be the species that initiates TLR4 signalling. These results indicate that as the disease progresses there are changes in the aggregate size distribution in CSF but not in the total number of species. This observation provides a method to follow disease progression that does not require the measurement of the absolute number of aggregate species. Moreover, we have also found a change in aggregate structure and the dominant mechanism of aggregate-induced cellular toxicity, when comparing MCI and AD CSF. Taken together, these results indicate that a combination of therapeutic agents targeting aggregated species with varying size and morphology, rather than a single agent targeting a single structure of one toxic form of Aβ may be needed to develop effective treatments for AD.

## Methods

### AD CSF

The CSF used for all the assays was collected by lumbar puncture from patients who sought medical advice because of memory problems. The samples were de-identified and aliquoted into 0.5 mL aliquots in polypropylene cryo tubes following centrifugation at 2,200 x*g* and stored at 80 °C pending experimental use. CSF Aβ1-42, T-tau and P-tau181 were quantified with sandwich ELISAs INNOTEST^®^ β-amyloid_1-42_, hTAU-Ag and Phospho-Tau [181P], respectively (Supplementary Table 1). All measurements were performed in clinical laboratory practice by board-certified laboratory technicians using procedures approved by the Swedish Board for Accreditation and Conformity Assessment. Intra-assay coefficients of variation were below 10%. All AD-positive samples had protein levels of Aβ42 < 600 ng/L, T-tau > 350 ng/L and P-tau181 > 80 ng/L. The study protocol was approved by the regional ethics committee at the University of Gothenburg.

### MCI and control CSF

MCI (6) and control (6) samples were collected from the Gothenburg H70 Birth Cohort Studies in Gothenburg, Sweden^14,36^. These samples were obtained from the Swedish Population Registry and included both persons living in private households and in residential care and approved by the Regional Ethical Review Board in Gothenburg. As described previously, lumbar punctures to collect CSF samples were performed in the L3/L4 or L4/L5 inter-space in the morning^37^. The first 10 mL of CSF were collected in a polypropylene tube and immediately transported to the laboratory for centrifugation at 1,800 x*g* in 20 °C for 10 min. The supernatant was gently mixed to avoid possible gradient effects, aliquoted in polypropylene tubes and stored at –70 °C^37^. CSF Aβ42, T-tau and P-tau181 were quantified with sandwich ELISAs INNOTEST^®^ β-amyloid_1-42_, hTAU-Ag and Phospho-Tau [181P], respectively.

As previously published, every 70-year-old in Gothenburg, Sweden, born during 1944 on prespecified birthdates was invited to the examination in 2014-2016, and 1203 participated (response rate 72.2%). Of these, 430 (35.8%) consented to a lumbar puncture^14^. Participants were examined at the Neuropsychiatric memory clinic at Sahlgrenska University Hospital in Gothenburg or at home. Experienced psychiatric research nurses performed the neuropsychiatric examinations, which comprised ratings of psychiatric symptoms and signs, tests of mental functioning, including assessments of episodic memory (short-term, long-term), aphasia, apraxia, agnosia, executive functioning and personality changes. Key informant interviews were performed by psychiatric research nurses as described previously^14,36^. Examinations included the Mini Mental State Examination and the clinical dementia rating (CDR). Dementia was diagnosed according to the DSM-III-R criteria.

For each participant, the clinical MCI diagnosis was determined in a clinical consensus conference, comprised of a neurologist and psychiatrist, taking into account the participants history and information from informants. CSF biomarker levels of Aβ 42, T-tau and P-tau were considered in choosing participants with MCI and underlying preclinical AD pathology. All MCI participants had a clinical dementia rating (CDR) score of 0.5. Healthy controls had to be free of MCI and Dementia and had a CDR of 0. Dementia diagnoses were an exclusion criterion in this study.

### Membrane permeabilisation assay

A detailed protocol and further description of the membrane permeabilisation can be found elsewhere^16^. Briefly, 200 nm mean diameter vesicles composed of 16:0-18:1 PC and 18:1-12:0 biotin PC (100:1) (Avanti Lipids) were prepared using extrusion and freeze-thaw cycles. Vesicles were filled with 100 µM of Cal-520 dye and tethered in the glass surface via biotin-neutravidin linkage. A glass surface was coated using PLL-g-PEG and PLL-g-PEG biotin (10:1) (Susos AG). To measure the background for each set, 9 different images were acquired in the presence of only 30 µL Ca^2+^ containing buffer, which we denote as blank (F_blank_). Then a 30 µL CSF aliquot was added to the glass coverslip and incubated for 15 min before images of the exact same fields of view were recorded (F_sample_). Then, 10 μL of ionomycin was added to the same coverslip and images of the vesicles were acquired in the same fields of view (F_ionomycin_). The recorded images were analysed to determine the fluorescence intensity of each vesicle under the three different conditions and the average Ca^2+^ influx for each vesicle was calculated using the formula (F_sample_ - F_blank_) / (F_ionomycin_ - F_blank_) X 100%. The stage movements were performed using a bean shell-based program which allows to select fields of view without any user bias. For antibody experiment, CSF and 300 nM antibody were incubated for 15 minutes and added to the vesicle containing glass coverslip for membrane permeabilisation study.

Imaging of individual vesicles were performed using a home-built total internal reflection fluorescence (TIRF) microscope. For excitation, a 488 nm laser (Toptica) beam was focussed in the back focal plane of the 60x, 1.49 NA ooil-immersion objective lens (APON60XO TIRF, Olympus, N2709400). Fluorescence emission from the dyes were collected by the same objective and imaged onto an air-cooled EmCCD camera (Photometrics Evolve, EVO-512-M-FW-16-AC-110)

### Inflammation assay

The BV2□cell line was derived from immortalised murine neonatal microglia and grown in 10% foetal bovine serum and 1% L-Glutamine supplemented Dulbecco’s Modified Eagle’s (DMEM) medium. The cells were incubated at 37□°C in a humidified atmosphere of 5% CO2 and 95% air, until the cell density reached approximately 1.6 x 10^6^□cell/mL. 200 µL of CSF was diluted in 1 mL of DMEM and added to BV2 microglia cells. Every 24 h the supernatant was removed for analysis and replaced by fresh diluted CSF. The TNF-α concentration in the supernatant was quantified using the Duoset® enzyme-linked immunosorbent assay (ELISA) development system (R&D Systems, Abingdon, Oxfordshire, UK). Three wells for each CSF were used to estimate variation in the experiments. For antibody experiment, CSF and antibody were diluted in DMEM buffer and added to the vesicle containing glass coverslip.

### Aptamer-DNA PAINT (AD PAINT) imaging

AD PAINT was performed as described previously^22^. Briefly, glass-slides were cleaned with argon plasma, rinsed with 1% tween-20 and washed with PBS. All buffers were first passed through a 0.02 µm filter (Anotop25, Whatman). The CSF was diluted ten-fold in PBS and added to wells on the coverslip formed by a multiwell chamber coverslip (CultureWell CWCS-50R-1.0). After 5 min the CSF was removed, the wells were washed with PBS and then filled with imaging mix (1 nM imaging strand (sequence CCAGATGTAT-CY3B) and 100 nM aptamer-docking strand (sequence GCCTGTGGTGTTGGGGCGGGTGCGTTATACATCTA) in PBS). The wells were then sealed using another clean coverslip. The samples were imaged on a home-built TIRF microscope using a 1.49 N.A., 60x TIRF objective (UPLSAPO, 60X, TIRF, Olympus) mounted on a Ti-E Eclipse microscope (Nikon) fitted with a perfect focus system. The Cy3B was excited at 561 nm (Cobalt Jive, Cobalt) passed through a FF01-561/14-25 excitation filter (Semrock). Fluorescence was separated from the excitation light using a dichroic mirror (Di01-R405/488/561/635, Semrock), passed through a filter (LP02-568RS-25, Semrock) and focussed on the EMCCD camera described above operating in frame transfer mode (electron-multiplying Gain of 11.5 e-1/ADU and 250 ADU/photon). To eliminate user bias an automated script (Micro-Manager^38^) was used to collect images in a grid. 5000 frames were collected with an exposure time of 50 ms. Images were analysed using the Peak Fit ImageJ plugin of the GDSC Single Molecule Light Microscopy package and custom scripts written in Python. The data analysis is described in detail in Whiten *et al*^28^.

### Atomic force microscopy imaging

CSFs are diluted 10x using PBS buffer and image on freshly cleaved mica substrates using AFM. 10 μL diluted CSF samples were deposited on the substrate at room temperature. The samples were incubated for 10 min and followed by rinsing with 1 mL milli Q water. Then the samples were dried using a gentle flow of nitrogen gas. AFM maps were created using a NX10 (Park Systems, city, South Korea) and JPK nanowizard2 system (JPK Instruments, city, Germany) operating in tapping mode. This set-up equipped with a silicon tip with a nominal radius of <10 nm. Scanning Probe Image Processor (SPIP) (Image Metrology, city, Denmark) software were used for image flattening and single aggregate statistical analysis. The lateral resolution of this technique is determined by the geometry of the AFM tip, although the measurement of the height of individual species is not significantly affect by the tip geometry. The average level of noise for each image was measured using SPIP software and is well below 0.1 nm. The signal to noise ratio for measuring a single Aβ42 monomer is close to 10. Thus, the characterisation of cross-sectional height can be performed with high sensitivity and accuracy.

### Confocal experiments using pFTAA

Single-molecule confocal experiments were performed using the previously-reported method^39^. CSF samples were diluted 1:1 into pFTAA solution (60 µM, PBS) and withdrawn through a single-channel microfluidic device at a flow velocity of 0.56 cm·s^-1^. A 488 nm laser beam (1.5~mW, Spectra Physics Cyan CDRH) was directed to the back aperture of an inverted microscope (Nikon Eclipse TE2000-U). The beam was reflected by a dichroic mirror (51008BS, Chrome) and focussed to a concentric diffraction-limited spot, 10 µm into the solutions in the microfluidic channel through a high numerical aperture oil immersion objective (Apochromat 60X, NA 1.40, Nikon). Fluorescence was collected using the same objective, passing through the same dichroic mirror and imaged onto a 50 µm pinhole (Melles Griot) to remove out-of-focus light. The emission was filtered (535AF45, Omega) and directed to an avalanche photodiode (APD, SPCM-14, Perkin-Elmer Optoelectronics). A custom-programmed field-programmable gate array, FPGA (Colexica), was used to count the signals from the APD and combine these into time-bins which were selected according to the expected residence time of molecules passing through the confocal probe volume. At each time-point data were collected for 10 minutes (100000-time bins, bin-width 0.2~ms). The experimental output data were collected using an FPGA card and analysed in Python using custom-written code. A threshold of background + 20 was set for all measurements so as to maximise the number of events, whilst removing the noise.

### Production and purification of Nb3 nanobody and designed antibodies

The Aβ-specific nanobody Nb3 was isolated from a llama (*Llama glama*) and amplified from the peripheral blood lymphocytes, as described previously^19^. The concentration was estimated by absorbance spectroscopy at 280□nM using a molecular extinction coefficient, which was calculated based on the sequence of the protein of 21,555□M ^1^ cm ^1^.

Rationally designed antibodies were generated as previously described^24^. Briefly, complementary peptides were selected using the cascade method to target linear epitopes within Aβ42 that scan its entire sequence. These complementary peptides were then grafted into the complementarity determining region 3 of an antibody scaffold by means of a mutagenic polymerase chain reaction with phosphorylated oligonucleotides. Protein concentration was determined by absorbance measurement at 280 nm using theoretical extinction coefficients.

### Statistical tests

To assess the statistical significance of the difference among AD, MCI and control CSF for membrane permeability assay, we performed a two-sample t-test (unpaired) (**Fig. 1b**) using origin 9.0. We also used the same two-sample t-test (unpaired) assay to check if antibody can significantly inhibit CSF-induced membrane permeabilisation (n=3) (**Fig 2b**. and **Supplementary Fig. 2**). To determine the statistical significance of the results on the inflammatory response induced by AD, control and MCI CSF samples (**Fig. 1d**) and on the antibody-induced inhibition of the toxicity of AD CSF samples we also performed two-sample t-test (unpaired) at 96 h (n=3) (**Fig. 2a**). The number of aggregates present in the CSF samples was tested using a one-way ANOVA with Tukey’s Multiple Comparison post-test (**Fig. 3b** and **Supplementary Fig. 1**). Significance in the differences of the size distributions of the aggregates were evaluated using the Kolmogorov-Smirnov test (**Fig. 3f, g**).

## Data availability

All data and codes are available from the authors upon reasonable request.

## Author Contribution

SD. DRW, CH, FSR, MR, DIS and CGT performed experiments and analysed the data. KB, IS, SK and HZ collected, analysed and provided the CSF samples. SM provided Nb3 antibody and FAA and MV provided the rationally designed antibody. SD, DRW, MV and DK wrote the manuscript. All authors discussed the results and contributed to the manuscript writing. SD and DK conceived the idea, designed the study. DK supervised the project.

### Acknowledgement

D.R.W. is supported by a Herchel Smith Fellowship. C.G.T. is supported by the European Research Council (669237), KB is supported by the Torsten Söderberg Foundation, Sweden. F. A. A. is supported by a Senior Research Fellowship award from the Alzheimer’s Society, UK. SK was supported by the Swedish state under the agreement between the Swedish Government and the county councils, the ALF-agreement (ALFGBG-813921, ALFGBG-65930, ALF-GBG-716681) and the Swedish Alzheimer Foundation. The Gothenburg H70 Birth Cohort Study was supported by grants from the Swedish Research Council (grants no 2015-02830, 2013-8717), Swedish Research Council for Health, Working Life and Wellfare (grants no 2013-1202, 2013-2300, 2013-2496), Alzheimerfonden, Hjärnfonden and the Swedish state under the agreement between the Swedish government and the county councils, the ALF-agreement (grants no ALF 716681).HZ is a Wallenberg Academy Fellow supported by grants from the Swedish Research Council (grants no 2018-02532), the European Research Council (grants no 681712) and Swedish State Support for Clinical Research (ALFGBG-720931). DK was supported by the Royal Society, the European Research Council with an ERC Advanced Grant (grants no 669237) and ARUK.

