## Supplementary Table 1 for "ADSoluble aggregates present in cerebrospinal fluid change in size and mechanism of toxicity during Alzheimer’s disease progression"

|  | Sample No | Total Tau ng/L | Aβ ng/L | P -tau ng/L | Age | Gender |
| --- | --- | --- | --- | --- | --- | --- |
| **AD** | 1 | 1400 | 270 | 155 | 74 | F |
|  | 2 | 769 | 348 | 94 | 59 | F |
|  | 3 | 763 | 302 | 86 | 81 | M |
|  | 4 | 836 | 507 | 89 | 69 | F |
|  | 5 | 1090 | 492 | 109 | 69 | F |
|  | 6 | 713 | 518 | 84 | 79 | F |
|  | 7 | 1030 | 540 | 197 | 66 | M |
|  | 8 | 1290 | 447 | 203 | 70 | F |
|  | 9 | 597 | 581 | 85 | 73 | M |
|  | 10 | 1010 | 552 | 92 | 82 | F |
| **MCI** | 11 | 634 | 436 | 81 | 70 | F |
|  | 12 | 554 | 258 | 73 | 70 | F |
|  | 13 | 362 | 463 | 50 | 70 | M |
|  | 14 | 354 | 295 | 67 | 70 | M |
|  | 15 | 566 | 371 | 78 | 70 | M |
|  | 16 | 384 | 345 | 49 | 70 | F |
| **Control** | 17 | 231 | 728 | 36 | 70 | F |
|  | 18 | 265 | 838 | 38 | 70 | F |
|  | 19 | 205 | 724 | 36 | 70 | F |
|  | 20 | 265 | 762 | 44 | 70 | F |
|  | 21 | 286 | 1000 | 49 | 70 | M |
|  | 22 | 252 | 792 | 36 | 70 | M |

**Supplementary Table 1. Characterisation of the CSF samples used in this study.**

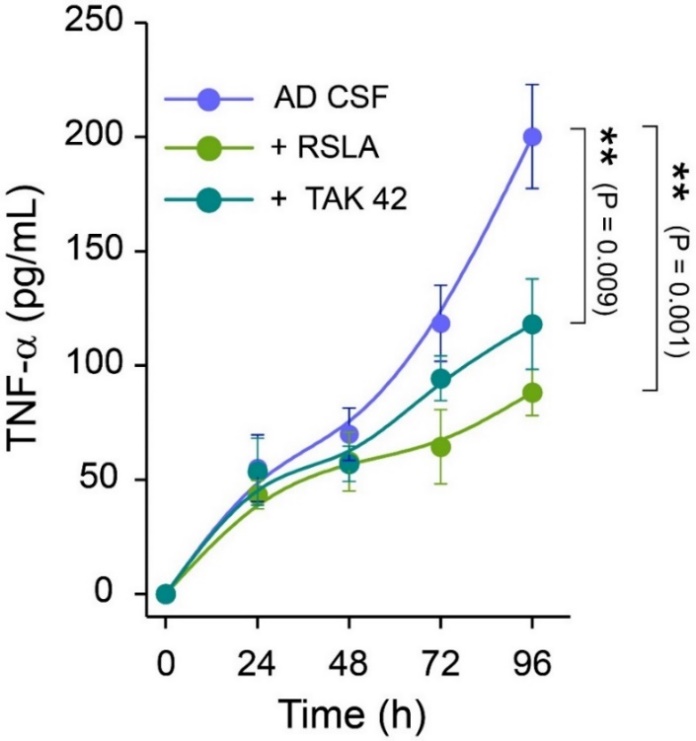

**Supplementary figure 1: Toll like receptor 4 (TLR4) antagonists block AD CSF-induced aggregate induced inflammation.** TAK-242, a small molecule inhibitor of TLR-4 and a known TLR4 antagonist *Rhodobacter sphaeroides* lipid A (RSLA) inhibits the AD CSF-induced inflammatory response by selectively binding to TLR4 and disrupting the interactions of TLR4 with its adaptor molecules (n=3, error bars is standard deviation). One way annova followed by post-hoc turkey were performed to compare the data sets

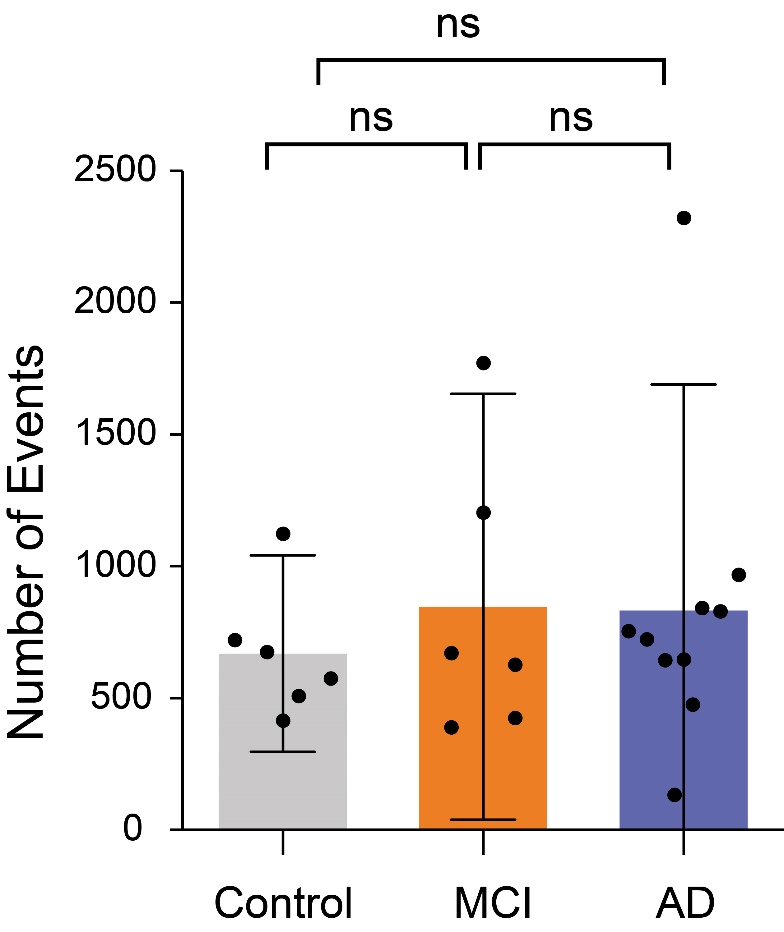

**Supplementary figure 2. Detection of aggregates present in control, MCI and AD CSF using pFTAA.** Pentameric formylthiophene acetic acid (pFTAA) is known to bind amyloid aggregates with high affinity. There is no significant difference in the number of pFTAA-active species in control, MCI and AD CSF. One way annova followed by post-hoc turkey were performed to compare the data sets
